# Engineering Principles and Algorithmic Design Synthesis for Ultracompact Bio-Hybrid Perfusion Chip

**DOI:** 10.1101/2022.03.16.484492

**Authors:** Amelie Erben, Thomas Kellerer, Josefine Lissner, Constanze Eulenkamp, Thomas Hellerer, Hauke-Clausen-Schaumann, Stefanie Sudhop, Michael Heymann

**Affiliations:** Center for Applied Tissue Engineering and Regenerative Medicine, Munich University of Applied Sciences, Lothstr. 34, 80533 Munich, Germany; Heinz-Nixdorf-Chair of Biomedical Electronics, TranslaTUM, Campus Klinikum rechts der Isar Technical University of Munich, Einsteinstraße 25, 81675 Munich, Germany; Center for NanoScience (CeNS), Ludwig-Maximilians-University, Geschwister-Scholl Platz 1, 80539 Munich, Germany; Multiphoton Imaging Lab, Munich University of Applied Sciences, 80335 Munich, Germany; Hyperganic Group GmbH, Georgenstraße 38, 80799 Munich, Germany; Department of Applied Sciences and Mechatronics, Munich University of Applied Sciences, Lothstr. 34, 80335 Munich, Germany; Institute of Biomaterials and Biomolecular Systems, University of Stuttgart, Pfaffenwaldring 57, 70569 Stuttgart, Germany

## Abstract

Bioinspired 3D microfluidic systems that combine vascularization with extracellular matrix architectures of organotypic geometry, composition and biophysical traits can help advance our understanding of microorgan physiology. Here, two-photon stereolithography is adopted to fabricate freestanding perfusable 3D cell scaffolds with micrometer resolution from gelatin methacryloyl hydrogel derived from extracellular matrix protein. As a proof of concept, we introduce an ultracompact bio-hybrid chip layout to demonstrate perfusion and cell seeding of double-digit μm proteinaceous channels. This perfusion chip consists of a standardized microfluidic interface fabricated from standard resin and a GM10 bioink channel printed atop of this interface. In addition, we demonstrate that algorithmic design synthesis can recapitulate intact alveoli and capillary networks with tunable design parameters to implement vascularized alveolar tissue models. This approach will allow for a systematic investigation of cell-cell and tissue dynamics in response to defined structural, mechanical and bio-molecular cues and is ultimately scalable to fabricate organ-on-a-chip systems.

## Introduction

Organ-on-a-chip systems that mimic relevant aspects of human tissue physiology have emerged as powerful tools to study complex inter and intra tissue phenomena^[1],[2]^. Such microfluidic systems combine biological structures with perfusion to control and manipulate cellular microenvironments and to ensure sufficient nutrient supply for short and long term culturing^[3],[4]^. Prominent microfluidic materials, such as glass, PDMS, or thermoplastics, however, have only limited utility to realize the specific biomechanics and biochemistry required for complex cellular functions, such as growth, proliferation and differentiation. In addition, soft lithography and injection molding based 2D fabrication constrain the possible 3D geometrical conformance of organ-on-a-chip systems to their target tissue. 3D microfabrication and bioprinting of spatially defined protein matrices seek to overcome this bottleneck in organ-on-a-chip systems.

Additive manufacturing techniques are increasingly used to fabricate 3D microfluidics. However, despite their general design freedom, attainable 3D print resolution imposes practical limits^[5]^. Two-photon stereolithography (TPS) is one of the few techniques that can provide free-form structures at sub-micron accuracy, thereby enabling performance improvements to microfluidic applications^[6],[7]^. Previous TPS fabricated 3D microfluidics include nozzles and mixers for serial crystallography^[7],[8]^, in-chip scaffolds to study cell migration^[9]^, filters^[10],[11]^ and enzyme reactors^[12]^.

3D-printed biomimetic *in vitro* models combine microfluidic perfusion with organotypic 3D cell niches fabricated with natural hydrogels^[1],[13]^. By tuning hydrogel properties, the effect of selected mechanical and bio-chemical cues on cells can be investigated *in vitro* to further our understanding of differences between healthy and pathological conditions^[14]^. Hydrogel based liver^[15]–[17]^, heart^[18],[19]^, lung^[20]^ and kidney^[21]^ as well as complex organoids^[22],[23]^ without perfusion of a vascular network have already been realized^[13],[14],[24]^. Adding minute control of perfusion on the micro scale to such *in vitro* models can help to further our understanding of complex 3D tissue environments, such as the intricately branched lung alveoli. These cellularized sub-millimeter air filled sacs circumscribed by a less than 10 μm thin membrane mediate O_2_ and CO_2_ exchange with the vascular capillary network^[2],[25]^. Alveolar 3D structure, as well as their biophysical and biochemical properties are pivotal for respiration and to maintain lung health^[26]^.

Complex organ-on-a-chip architectures can be fabricated by stereolithography^[27]^, laser induced forward transfer (LIFT) ^[28]–[30]^, inkjet printing^[31]^ and extrusion based methods^[1],[15],[32]–[34]^. However, enhanced resolution at the sub-cellular level is required to ultimately recapitulate the complete 3D *in vivo* ultrastructure. While one-photon stereolithography and volumetric bioprinting achieve sufficient fabrication rates to generate clinically sized tissue structures with a two-digit μm scale resolution^[35],[36]^, TPS can further enhance the resolution of structured hydrogels down to the sub-micrometer scale^[37],[38]^. By initiating a two-photon absorption process within photo-activatable materials, complex 3D structures can be polymerized^[39]^. Such structures fabricated with synthetic resin materials have been shown to guide cell adhesion^[40]^ and differentiation^[41]^ when functionalized with cell-mediating bio-molecules. In addition, photoactivatable protein-based resins, can be used to realize biomimetic structures with the appropriate biochemical and biophysical cues^[37],[39],[42]^. Recent TPS based organ-on-chip devices printed with protein based resins include a biomimetic placental barrier model, separating the fetal from the maternal compartment^[43]^, as well as microvascular structures^[44]^. Nevertheless, both of these proteinaceous cell scaffolds were printed within predefined microfluidic channels limiting possible growth and expansion of the biological structures. Also, the proteinaceous microchannels were so far not individually interfaced to the microfluidic periphery, but fluid flow within channels was achieved by flushing the entire microfluidic encasement. Strategies to 3D-print free-standing proteinaceous micro-structures which can be readily and selectively interfaced to microfluidic devices and perfused, e.g. with cell culture medium or blood substitutes, are still missing today.

Another limitation, when it comes to 3D printed biomimetic structures with micrometer and sub-micrometer precision are computer aided design (CAD) programs. Existing CAD software is usually based on “manual” step-by step design principles intended and suitable for subtractive and formative manufacturing methods rather than organic designs for additive manufacturing. The resulting structures can hence deviate strongly from their natural tissue counterparts and small design changes of complex designs usually result in time consuming workflows^[45]–[48]^. Alternatively, tissue imaging dataset derived designs have been used^[37],[47],[49]^ to recapitulate native geometries accurately, but lack systematic variation and adjustment of individual design parameters^[45]^. Algorithmic design based on parametric and algorithmic modelling provides an alternative and allows to efficiently explore and optimize geometries based on a set of logical operations and user defined rules^[50]–[52]^. Algorithmic design and topology optimization algorithms can yield hierarchical branching patterns resembling those found in nature^[48],[53]^. Voronoi tessellation can mathematically describe for instance the alveoli through a unit cell geometry and a labyrinth connecting individual cells for ventilation^[25],[54],[55]^. Iterative algorithmic design approaches can emulate natural evolutionary strategies^[47]^ to achieve topology optimized designs with biomimetic appearances^[47],[50]^. Algorithmic design principles have already been successfully harnessed in architecture^[56],[57]^, aerospace^[58]–[60]^, medical implant^[61],[62]^ or bone scaffold^[48],[53],[55]^ design generation, and recently, simplified algorithmic models of alveolar tissue have been used to simulate their mechanical loading in computer modelling^[25]^. Algorithmic design of functional biomimetic shapes is hence also a promising avenue for organoid and organ bioprinting applications.

Here we present an ultracompact bio-hybrid chip with a freestanding perfuseable micro channel that was printed from ECM-derived protein. The chip contains a plastic base of about one mm^3^ in size that provides a robust fluidic connection between a nutrient media reservoir and the ‘micro-vasculature’ channel. Perfusion and cell seeding of the channel is demonstrated for an 80 μm diameter microchannel geometry fabricated from gelatin methacryloyl (GM10)^[63]^. In contrast to previous organ-on-a-chip devices, our configuration does not encapsulate the cultured biological system within a rigid casing, but rather allows it to potentially grow and expand beyond its initial confinement. Alveolar scaffolds of varying size, wall thickness and degree of vascularization were then designed using a generative algorithm and 3D-printed with two-photon stereolithography, demonstrating that a generative design together with high resolution 3D printing indeed allows to create complex biomimetic 3D structures. Taken together, this approach opens new avenues to design and 3D-print integrated organotypic 3D microtissues.

## 2. Results and Discussion

### 2.1 Ultracompact 3D bio-hybrid chip – design and assembly

To connect individual hydrogel scaffolds containing simple channels or complex, biomimetic microtissues, we first constructed and 3D-printed an adapter to seamlessly interface the hydrogel scaffolds with standard microfluidic equipment (Figure 1, SI Design 1). In a proof-of-principle experiment, a 400 × 215 × 160 μm GM10 gelatin methacryloyl^[37],[63]^ block (SI Design 2) containing an 80 μm diameter channel and a minimal wall thickness of 67.5 μm was printed onto the adapter. The channel was curved to position both in- and outlet at the interface between GM10 block and adapter at a distance of 70 μm. For the adapter we chose IP-S, a commercial acrylate based TPS resin. Channel sockets of the adapter measured 100 μm diameter to interface hydrogel caps and 368 μm diameter to receive 360 μm outer diameter glass capillary tubing^[7]^. Mounted glass capillaries where usually 10 cm long. Both the GM10 channel and the adapter were 3D printed via two-photon stereolithography using a 25x / 0.8 NA objective.

**Figure 1:**
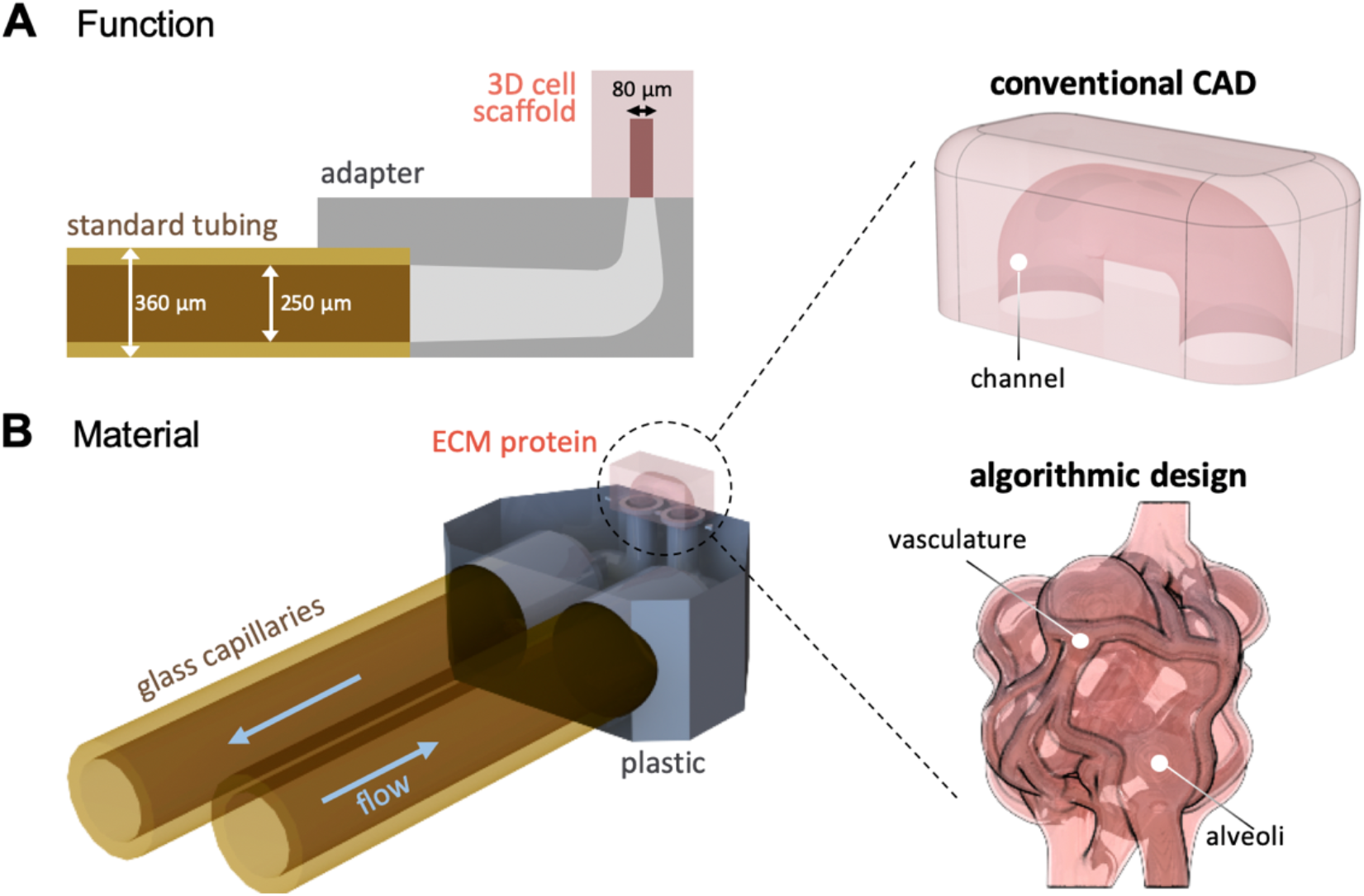
The bio-hybrid perfusion chip connects microscopic, precision 3D printed protein scaffolds with standard microfluidics. (**A**) A plastic adapter base (SI Design 1) bridges delicate channels embedded in proteinaceous 3D printed cell scaffolds (SI Design 2) with diameters of ∼80 μm to glass micro capillaries with inner and outer diameters of 250 and 360 μm, respectively. (**B**) As a proof of concept, we designed an adapter holding two capillaries for in- and outwards flow. Inside the adapter, the channel diameter is reduced to seamlessly interface with channels embedded in proteinaceous cell scaffolds such as simple channels generated using conventional CAD or algorithmic, biomimetic designs mimicking e.g. the alveoli synthesized using algorithmic design.

The complete adapter was larger than the field of view of the objective and hence printed as a sequence of 300 × 300 × 200 μm blocks (Figure 2A). Stitching interfaces between the blocks resulted in a rectangular line pattern on the surface, noticeable in scanning electron microscopy. To reduce printing time, the adapter was designed with minimal excess material and contained flat surfaces to limit lensing artifacts in microscopy imaging of internal channels. This also facilitated subsequent dip-in mode protein hydrogel scaffold printing on the adapter, which in theory can realize complex designs with arbitrary dimensions. The glass capillary ports of the adapter were placed in parallel to ease handling. L-shaped alignment features help orienting the hydrogel segment onto the adapter. After printing, the adapter units were developed in PGMEA solvent, rinsed in isopropanol, dried and then centered on 22 × 22 × 0.17 mm microscope cover glass slides and secured in position with polyimide adhesive tape (Figure 2B). Glass capillaries were then carefully inserted into the respective adapter ports under a stereomicroscope. Five-minute epoxy glue was mixed for one minute, further cured for another 30 seconds, and then applied to seal the capillaries into the adapter. Glue was applied dropwise using a capillary and allowed to cure for at least 10 minutes^[7]^.

**Figure 2:**
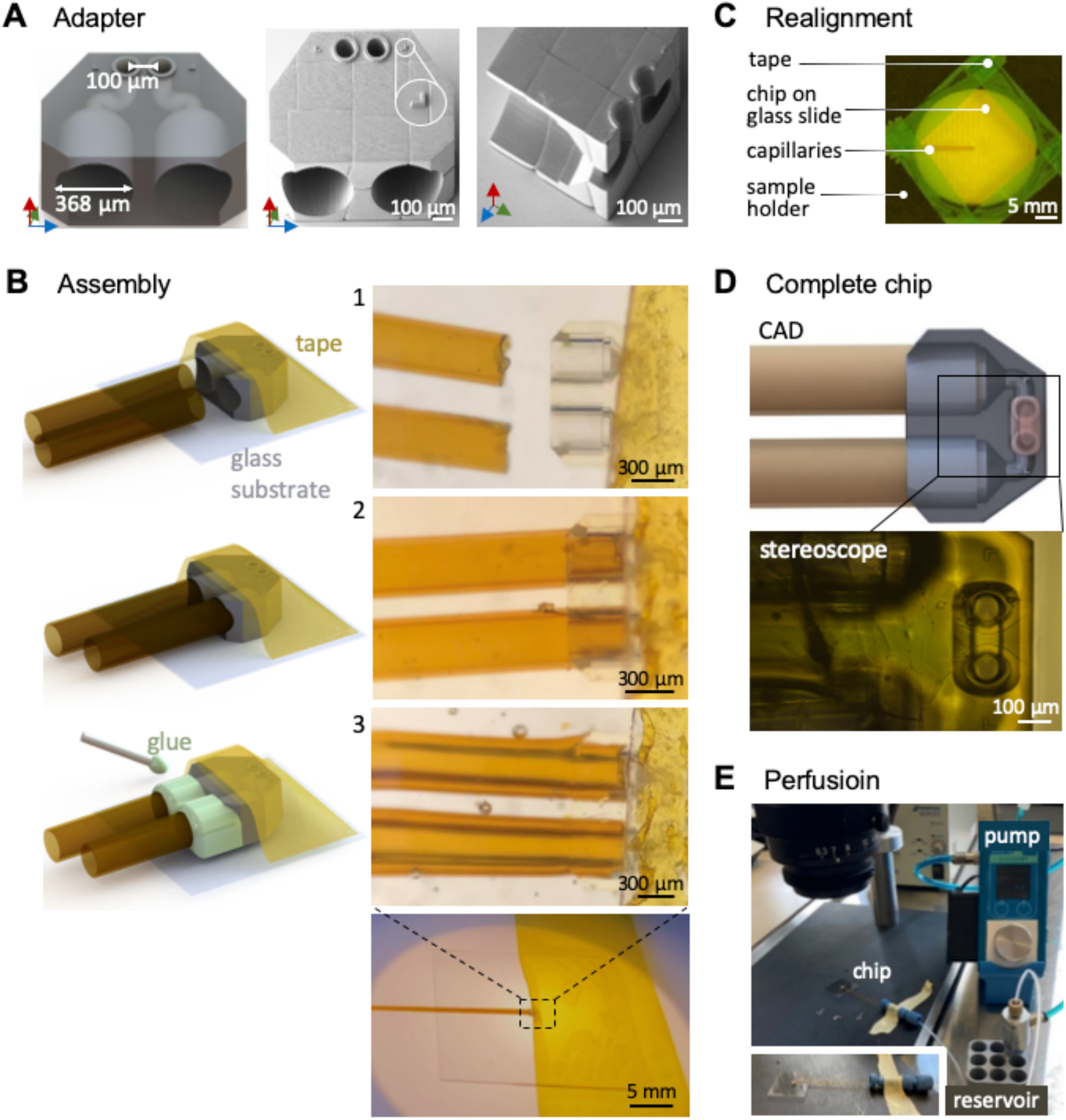
(A) The adapter interfaces hydrogel channels via 100 μm diameter channel ports and two standard 360 μm outer diameter HPLC capillaries via 368 μm diameter ports (left). Scanning electron microscopy image of entire (middle) and cut-open (right) adapters fabricated from IP-S show characteristic print block stitching lines, but otherwise unobstructed channels and ports. **(B)** After development, perfusion chips were secured to glass substrates using polyimide adhesive tape (1). Capillaries were inserted under a stereomicroscope (2) and epoxy glue was applied dropwise (3). After curing for 10 minutes, the adhesive tape was removed. **(C)** Glass slide holding an assembled chip with capillaries was reinserted and secured with adhesive tape on the two-photon stereolithography sample holder for **(D)** 3D printing of protein channels using GM10 resin. Schematic top view showing capillaries (brown), adapter (grey) and GM10 scaffold (pink) and corresponding micrograph of chip with GM10 scaffold. **(E)** Finished bio-hybrid perfusion chips were connected to a fluid reservoir and a pressure-controlled pump via standard fluid connectors.

In a final step, the glass slides holding the assembled adapters were reinserted into the TPS printer (Figure 2, C) by fixing them onto the sample holder using adhesive tape (green) and applying a drop of approximately 25 μL GM10 resin. The adapter unit was centered in the printer using L-shaped alignment marks to correctly place and seal the GM10 channel segment^[8]^. After printing, the GM10 channel was developed in PBS buffer and visually inspected in a stereomicroscope (Figure 2D). The completed bio-hybrid perfusion chip was then connected to a pressure-based flow control unit via the glass capillaries using MicroTight (IDEX Health & Science LLC) connectors and tubing for perfusion experiments (Figure 2E).

### 2.2 Shape fidelity and smallest continuous protein channel

Reference scaffolds were designed and printed to identify the lowest attainable channel diameter for the used GM10 TPS recipe. A hallmark of organ homeostasis is continuous oxygen and nutrient supply and waste material removal through branched vasculature. Distal microvasculature in human ranges between ∼5 – 30 μm in vessel diameter^[44],[64]^. To confirm suitable imaging conditions, blocks of 70 × 70 × 290 μm that enclosed an empty void with a 10 μm thick wall of polymerized GM10 protein resin were printed in petri-dishes and stored submerged in PBS to prevent drying. All scaffolds were imaged with a 20x / 0.95 NA objective using two-photon excited fluorescence microscopy to derive x/z cross sections from recorded z-stacks. To identify PBS content within the void, we imaged the scaffold submerged in 0.5 mg/mL FITC-CM-dextran solution (Fluoresceinisothiocyanat-Carboxymethyl-Dextran) with 150 kDa average molecular weight and a radius of gyration of order 8 nm^[65]^. The scaffold, which has been printed using a GM10 ink doped with 0.5 mg/ml rose bengal as a fluorescent stain was excited with a two-photon excitation wavelength of 1034 nm while FITC-CM-Dextran was exited at 780 nm (Figure 3A). The peak absorption and emission wavelengths were 559 nm and 571 nm for rose bengal, as well as 491 nm and 516 nm for FITC-CM-dextran. Hence emission filters to transmit between 610 - 900 nm for rose bengal and 468 - 552 nm for FITC-CM-Dextran were chosen. This configuration could image both fluorophores separately with minimal cross talk between both detection channels. Superimposed rose bengal / FITC images reveal only negligible FITC fluorescence within the closed void, confirming that FITC-CM-dextran solution cannot permeate the polymerized protein hydrogel (Figure 3B).

**Figure 3:**
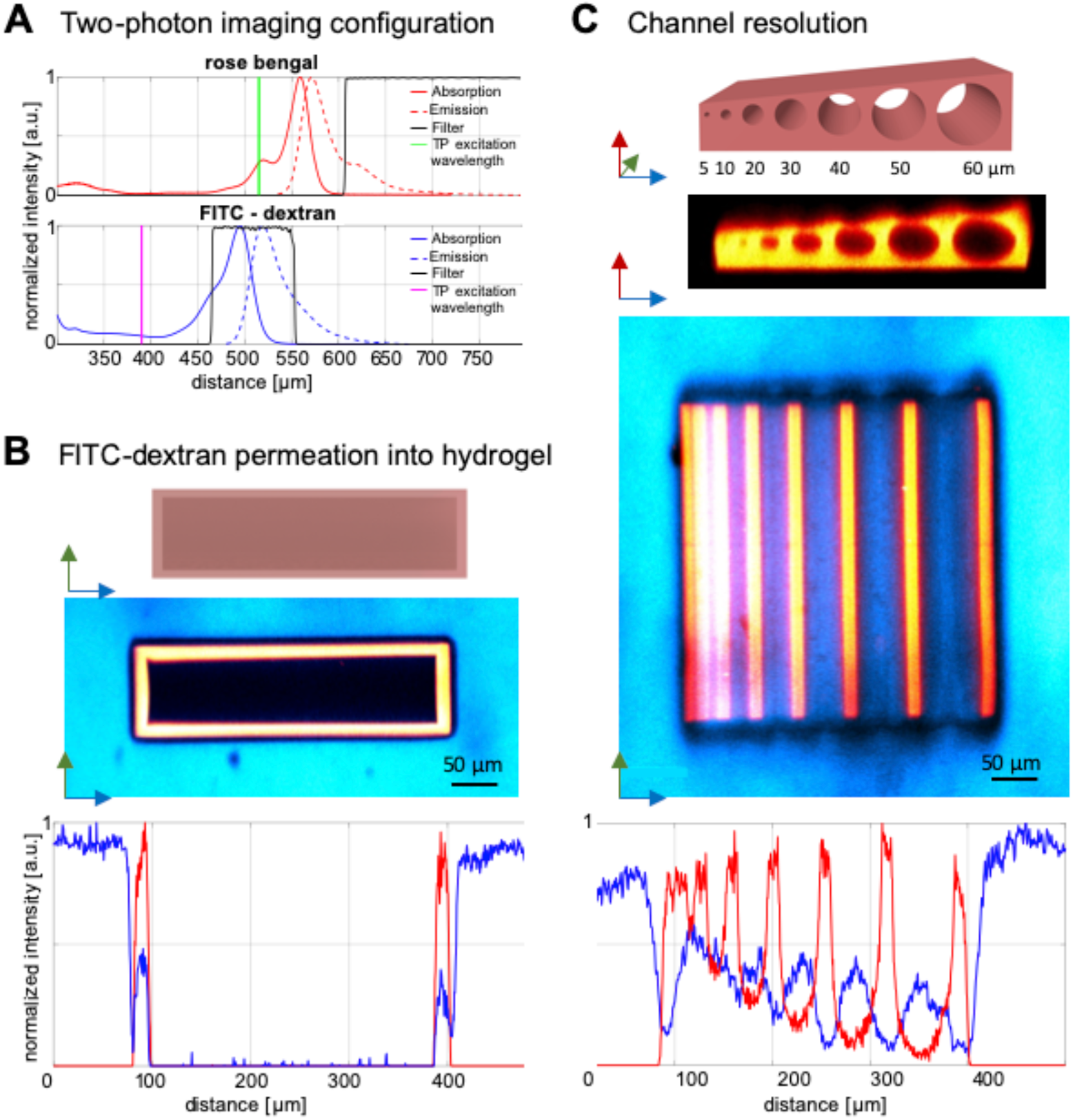
Assessing resolved microchannel diameters in 3D printed proteinaceous GM10 resin. **(A)** Two-photon microscopy imaging used 780 nm or 1034 nm two-photon excitation for aqueous FITC-CM-dextran (blue) and 0.5 mM rose bengal stained scaffolds (red) and matching 468-552 nm and 610-900 nm bandgap emission filters. PBS washing buffer was exchanged with 0.5 mg/mL FITC-CM-dextran to accessible internal channel sections after scaffolds were development via confocal sections and corresponding line plots **(B)** Large FITC-dextran of 150 kDa with a radius of gyration of order 8 nm, was not observed to permeate into the scaffold walls and central cavities remained non-fluorescent in both channels suggesting that the GM10 hydrogel mesh size was large enough to facilitate rose bengal efflux during scaffold development but small enough to prevent FITC-Dextran influx. **(C)** A test scaffold containing channels with diameters of 5 to 60 μm, each 290 μm long was designed and printed. A x/z section was derived from imaged z-stacks to compare the printed structure with the CAD model. Fluorescence image of the central the x/y plane (middle). The x/y section of both channels are overlaid normalized intensities along the cross-section indicating FITC-CM-Dextran permeated all channels down to 10 μm fully. Scale bars 50 μm

Next, for channel print-fidelity evaluation reference scaffolds were fabricated that contain hollow channels with diameters ranging from 5 to 60 μm (Figure 3C). Observed channel cross sections revealed non-spherical, nearly elliptical channels, which may be attributed to the fact that TPS and two-photon microscopy both have an elongated point spread function in z-direction, resulting in an increased polymerization and fluorescence detection volume along the z-axis. Fuzzy edges at the top of the structure with lower intensities compared to the bottom side most probably hails from absorption of the excitation laser, as it passes the scaffold from the bottom located at the glass slide surface to the scaffold top in the inverted microscope. Superimposed fluorescence images and intensity curves reveal FITC-CM-dextran in all channels from 60 μm to 10 μm. The 5 μm channel in turn was barely recognizable in the fluorescence images. Diffusion of FITC-CM-dextran into channels down to approx. 10 μm in diameter suggests that these channels can be perfused, rendering them suitable to cover a wide range of microvasculature dimensions in microfluidic devices. These results are in agreement with Dobos et al. who fabricated thiolated gelatin and gelatin-norborene channels down to 10 μm diameter using a comparable TPS configuration^[44]^. Nevertheless, to recapitulate also the smallest entities of the native microvasculature, a future reduction of channel diameters to below 10 μm seems desirable^[66]^. This may be achieved by using higher numerical aperture writing objectives as well as suitable photoabsorbing resin additives such as tartrazine ^[35]^ to further increase print resolution.

### 2.3 Active perfusion and cell seeding in ultracompact bio-hybrid chip

After evaluating shape fidelity and printing resolution, we now focus on liquid transport through the channels as well as the possibility of seeding cells within the channels. First, we demonstrate perfusion of an individual 80 μm diameter hydrogel channel, which is convenient to fabricate and large enough to avoid the risk of clogging with cells. Perfusion was tested by pushing PBS liquid from a reservoir into a connected chip using 345 mbar pressure (Figure 4, Movie S1). Inflowing PBS buffer successively removed air bubbles from the channel until the entire system was bubble free after a few seconds. The evacuation of air bubbles confirmed a water tight seal between the IP-S contact chip and the protein channel, implying sufficient adhesion and precise fitting between both materials for the applied pressure gradient. In the future, perfusion of smaller diameters will be tested by decreasing the port dimensions on the adapter. To mimic interveined microvasculature, the complexity of perfusion chips can be enhanced to entail several channels, which can be individually contacted. Hierarchically branched vasculature may be recapitulated within protein scaffolds starting from the 80 μm inlets and gradually decreasing protein channel diameters.

**Figure 4:**
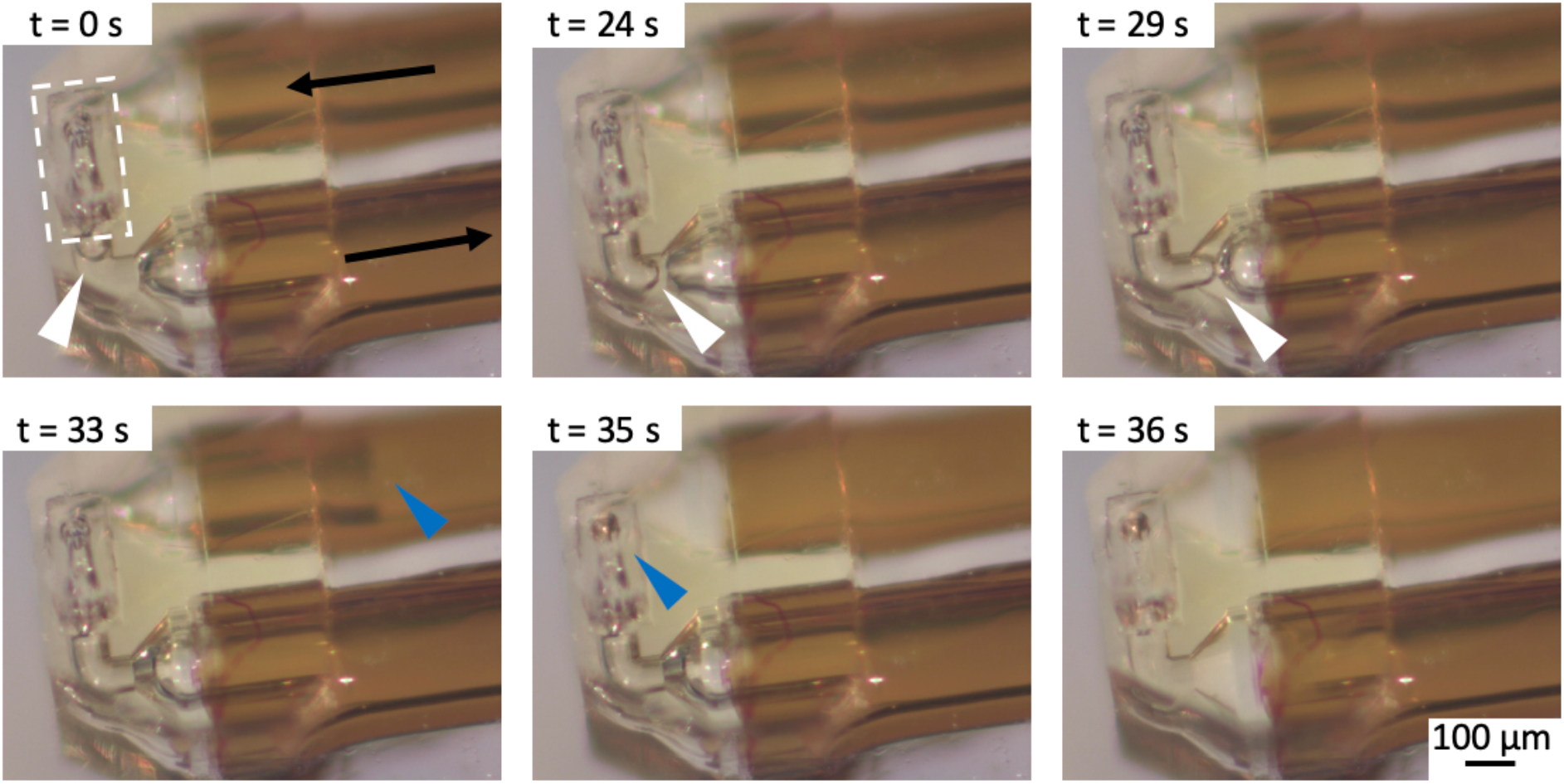
Liquid flow in bio-hybrid perfusion chip. The adapter chip is submerged in PBS buffer to prevent drying induced damage to the proteinaceous GM10 hydrogel cap (white dashed line at t = 0s) while PBS buffer and air are flushed through it (Movie S1). Black arrows indicate flow direction, while white/blue arrows show the advancing air/PBS buffer meniscus. An air bubble was pinned inside the adapter chip (white arrows), but was flushed out when a continuous water column (blue arrows) progresses from the inlet capillary through the channel (t = 33-35 s) until the chip is air bubble free after 36 s.

In the next step, assembled bio-hybrid chips were seeded with GFP labelled human mesenchymal stem cells (hMSCs) of the SCP1 cell line^[67]^, a comparably robust cell line. Chips were connected to a pressure controlled pump to load hMSCs together with cell type specific culture media into the 80 μm diameter GM10 channel segment. Combined bright field and epifluorescence microscopy was used to confirm cell seeding inside the GM10 channel. However, high auto fluorescence of the 3D printed, acrylate based adapter (Figure 5, left) required thresholding and background subtraction to resolve individual cells within the GM10 channel (Figure 5, right). While cells could be introduced into the 3D printed proteinaceous channel without clogging, future experiments should investigate cell adhesion to the hydrogel channel walls during long-term cultivation under perfusion. Especially the response of epithelial cells native to the capillaries will be interesting to access in this regard. Also, lower auto-fluorescent resins could be advantageous for specific application scenarios^[68]^.

**Figure 5:**
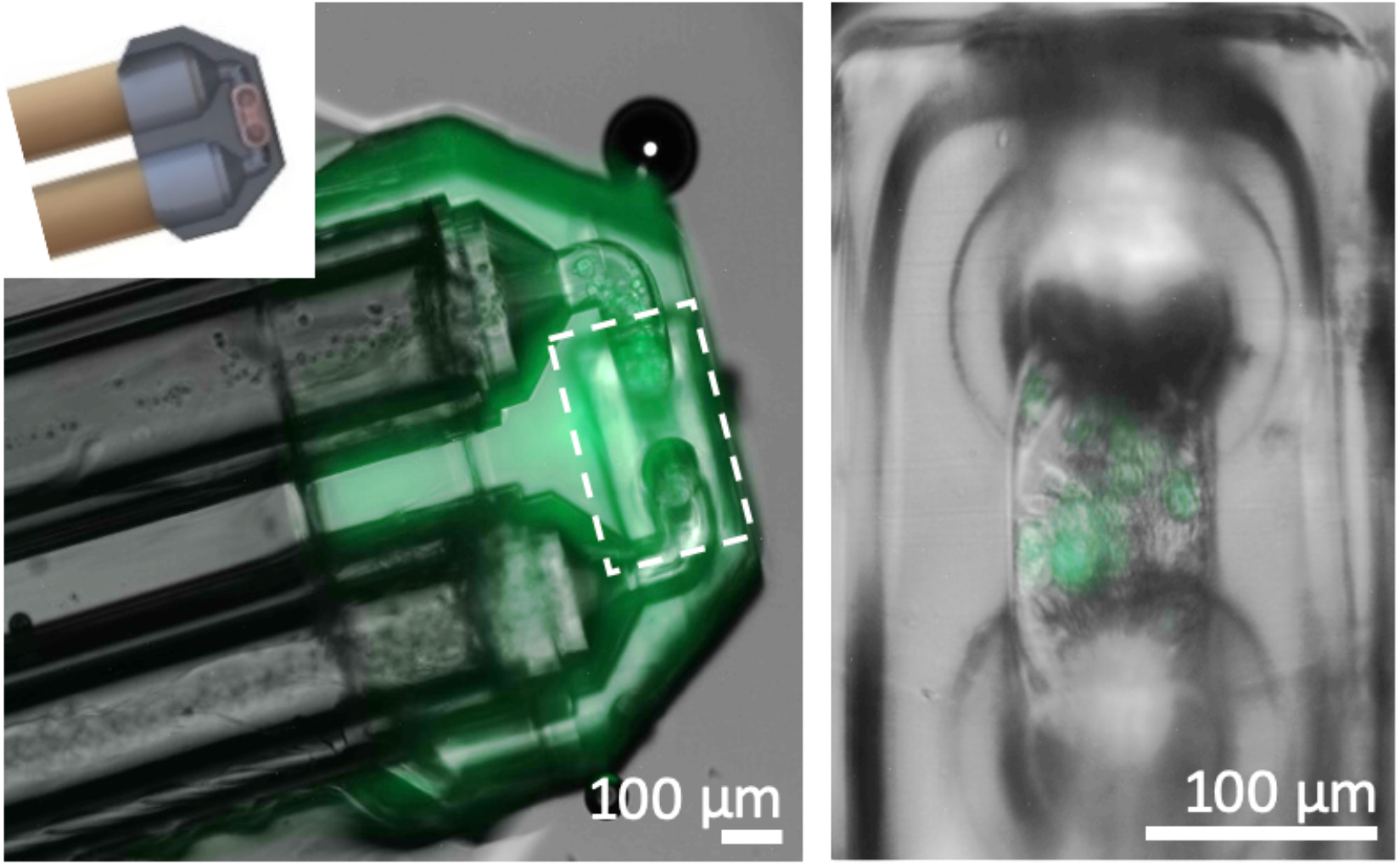
Bio-hybrid perfusion chip colonized with human mesenchymal stem cells that constitutively express GFP. Cell suspension (left) was introduced via the glass capillaries into the GM10 channel highlighted by a dashed white rectangle. A close up reveals green fluorescent hMSCs within the GM10 channel segment (right).

### 2.4 Reusability of assembled adapters and hydrogel biodegradability

Recycling of sub-components can reduce the fabrication effort for multi-step assemblies. To evaluate the reusability of the adapter chip, the hydrogel block was carefully stripped away from the adapter using a plastic pipette, followed by thoroughly rinsing with ddH_2_0. The glass capillaries for liquid supply were left intact in the adapter. Electron microscopy of a stripped adapter revealed only minor gelatin residuals remaining on the adapter surface as well as a protein gel thread inside one of the two hydrogel contact ports (Figure 6). Simple mechanical cleaning can hence suffice to reuse adapters and may be complemented by enzymatic degradation of the biomaterial. Ultrasonic cleaning or washing with sodium hydroxide compromised the integrity of the glued glass capillary seal.

**Figure 6:**
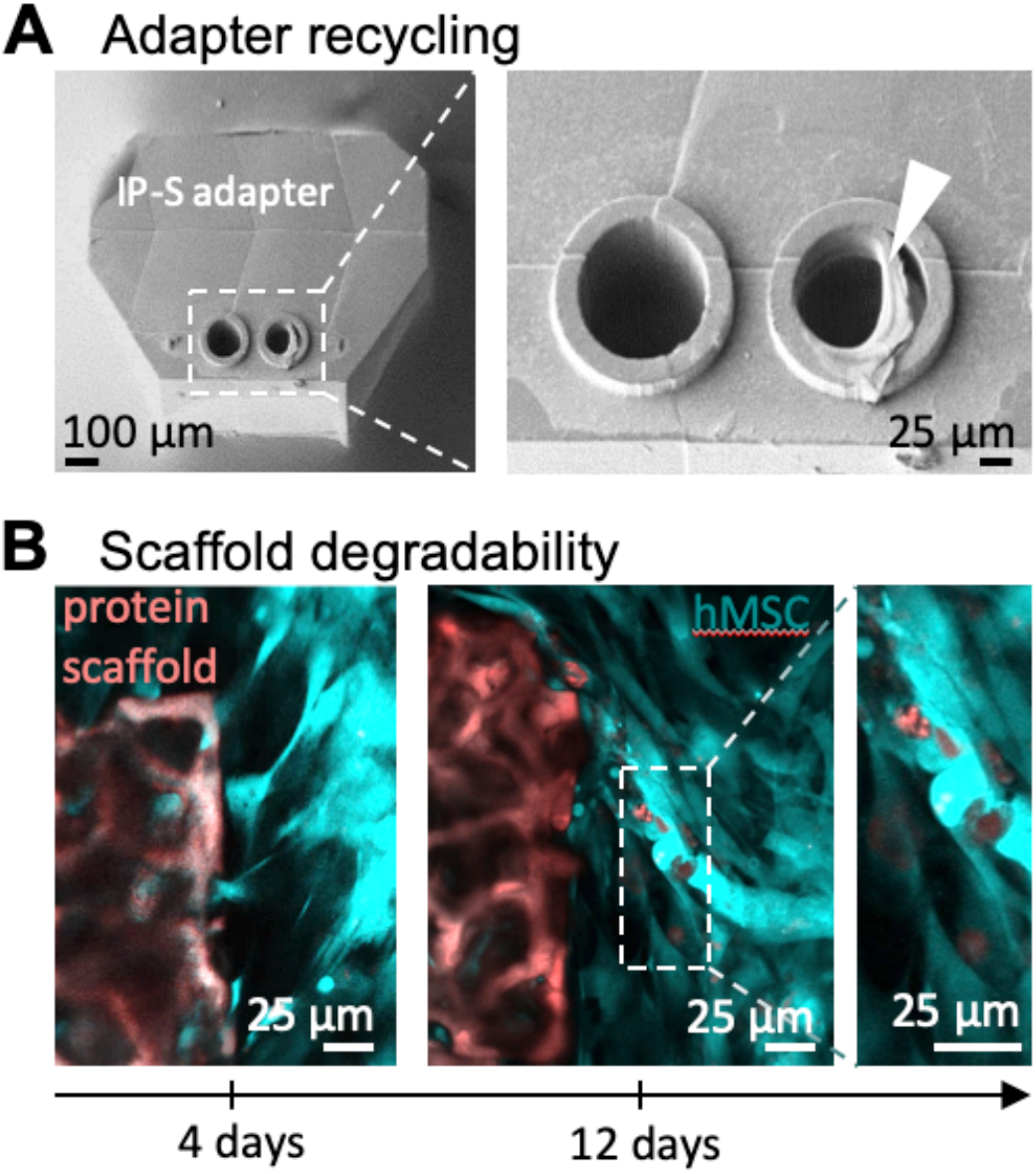
Protein hydrogel chip turnover. **(A)** Scanning electron microscopy images of adapter unit after mechanically removing GM10 block. A close up reveals a residual GM10 gel thread within one of the two adapter ports (white arrow). **(B)** Protein scaffolds (red) 4 and 12 days after colonization with human mesenchymal stem cells (hMSC, blue). After 12 days, cells have gradually degraded the scaffold.

For many applications, defined biodegradability of proteinaceous scaffolds is important. Ideally, artificial scaffolds can provide shape and mechanical strength until cells have secreted and modeled their own natural ECM, which then replaces the artificial material. Figure 6B shows GM10 scaffold degradation by hMSCs (SCP1 cell line) over the course of 12 days after cell seeding. While after 4 days, the scaffold still seems to be mostly intact, cells are able to remove small chunks from the scaffold by day 12, demonstrating that the cross-linked GM10 can be resorbed by the hMSCs. This is in agreement with Van Hoorick et al., who demonstrating enzymatic biodegradation of cross-linked gelatin scaffolds^[69]^.

### 2.5 Biomimetic design synthesis of perfusable protein scaffolds

Alveoli are air filled sacs that exchange gas with a highly branched microvasculature at the distal ends of the lung^[70],[71]^. A thin membrane lined with pneumocytes regulates O_2_ and CO_2_ exchange between red blood cells and the gas phase (Figure 7A)^[2],[72]^. Alveoli diameters range from approx. 58 μm in mice^[73]^ to approx. 200 μm in humans^[74]^. In a previous study, we cellularized precision 3D printed scaffold geometries, derived by imaging native alveoli tissue^[37]^. This imaging based approach failed however to generate intact perfusable capillaries within these alveoli tissue templates. In addition, the design could not be adapted to varying shapes and sizes, such as wall thickness or alveoli size.

**Figure 7:**
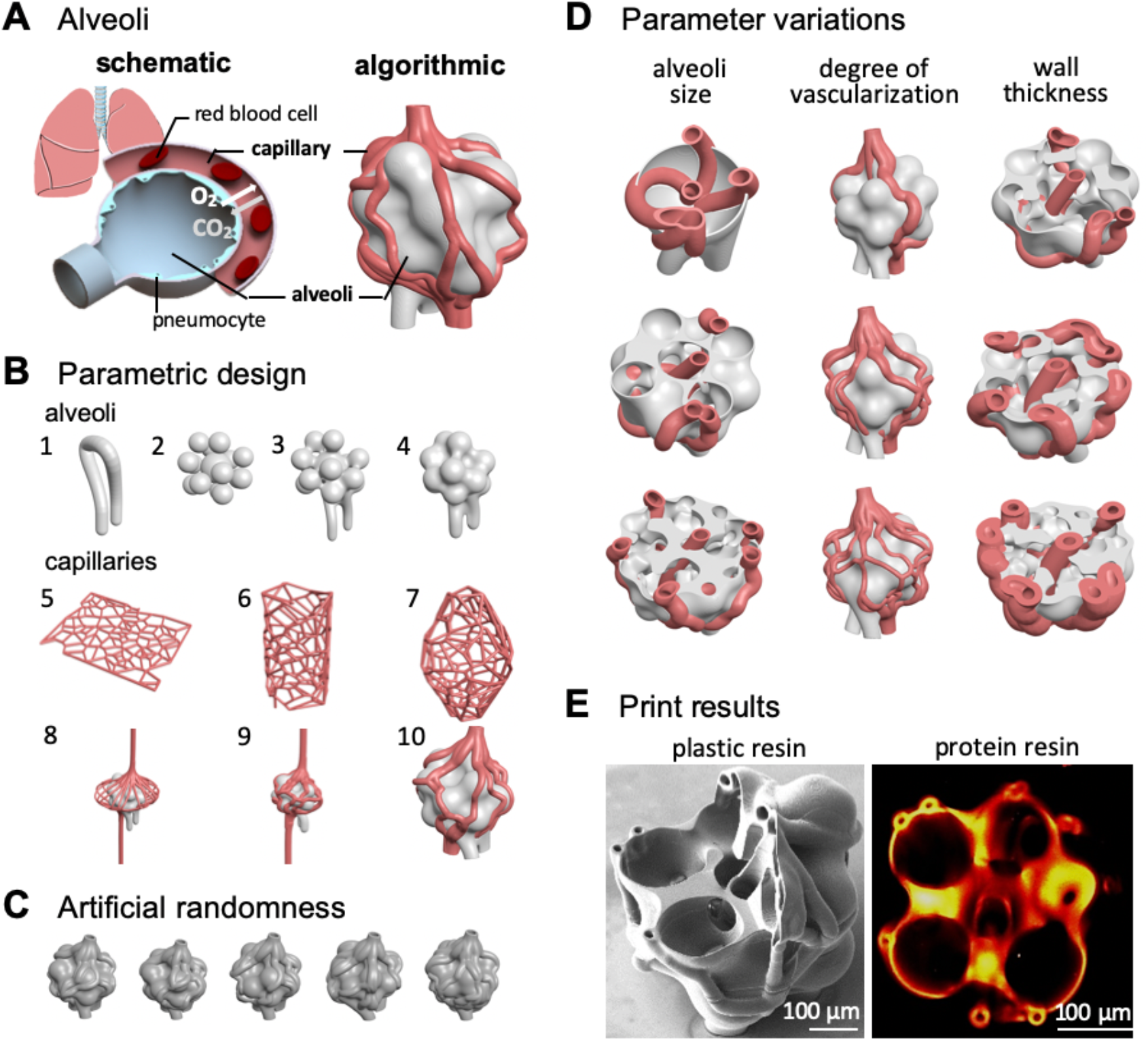
Parametric design synthesis of vascularized alveoli. **(A)** Lung alveoli are air-filled sacs lined with pneumocytes to mediate CO_2_/O_2_ exchange. Algorithmic design can generate alveoli models (right) with biomimetic capillary networks (red) and interconnected air sacs (grey). **(B)** Algorithmic design computation proceeds in several stages: (1) The air path is defined through lattice beams. (2) A spherical foam map mimics alveolar sacs of pre-defined min./max. sphere sizes and gaps. (3) Sphere surfaces are connected by introducing offsets from individual spheres, before the air path is joined with the spherical foam map. (4) The external surface is then smoothed. (5) Starting from polygonal cells based on random input points on a 2D canvas, (6) a transformation to a 3D cylindrical assembly (7) is followed by defining variable radii, a spine, height variable beam thickness and edge curving. (8) This cylindrical Voronoi mesh is then partially snapped onto the previously designed alveolar base (9) using transition interpolation. (10) Final in- and outlets are trimmed. **(C)** Design features such as sphere layout or capillary paths can be randomized to select degrees. **(D)** Biomimetic algorithmic designs (Movie S2) allow to explore specific design parameter variations (Figure S1), such as alveoli size, degree of vascularization and wall thickness to systematically investigate single cell and tissue dynamics in response to context specific 3D design cues. **(E)** Parametrized alveoli templates were printed true-to-scale using two-photon stereolithography in both standard acrylic-based resin (scanning electron microscopy, left) and proteinaceous GM10 resin (Movie S3, two photon fluorescence microscopy, right).

To experimentally realize perfusable biomimetic microtissue, we now designed an alveoli network using algorithmic design. Interconnected alveoli design synthesis (Figure 7B) was initiated by defining an air path using lattice beams^[58]^, followed by a spherical foam map defined to mimic the alveoli through a set of minimum and maximum sphere radii and gap dimensions between individual spheres within a bounding geometry. Multiple spheres were joint by defining an offset from each spherical surface and connected with the initial air path contour. This was followed by a sequence of offset and smoothing commands to hollow the geometry and to achieve the final alveolar configuration. Design synthesis of the bounding capillary network proceeded through creating polygonal cells seeded from random input points on a 2D canvas that were derived using a Voronoi algorithm, followed by conversion to a 3D cylindrical arrangement and integration of a variable radius, a spine and variable beam thickness along the height, as well as curving of edges. Subsequently, the cylindrical Voronoi mesh was partially snapped onto the previously designed alveolar base and interpolated, before trimming in- and outlets to their final lengths. Resulting alveoli were hollow and surrounded by a capillary network. Both alveoli and capillaries can be contacted through distinct in- and outlets for cell seeding, medium perfusion and tidal ventilation. In addition, all desired alveoli in-/outlets are located in one plane to interface matching ports of the IP-S adapter chip (Movie S2).

Contrary to conventional CAD, our algorithm easily accommodates artificial randomness at all design stages (Figure 7C), as each parameter may be set to a defined value or left to permutate within the defined constraints. Here we left total construct size and air path constant, while capillary paths and individual sphere locations were randomly distributed. The resulting geometries hence vary slightly in their design along defined parameters. In addition, this algorithmic design approach allows for deliberate design permutations such as alveoli size, degree of vascularization and wall thickness (Figure 7D, Figure S1).

Using dip-in mode TPS printing, the computational design was printed both with acrylate-based IP-S resin and imaged with scanning electron microscopy (Figure 7E, left), as well as with gelatin-based GM10 resin and imaged with two-photon fluorescence microscopy (Figure 7E, right, Movie S3). Our approach of designing a biomimetic alveoli model includes features to recapitulate breathing dynamics and controlled medium distribution in accordance with the bioinspired alveolar models from Grigoryan et al.^[35]^ with diameters of ∼1 mm for alveoli and ∼300 μm for vasculature realized in single photon stereolithography. Our higher resolution TPS process achieve diameters ranging from 80-170 μm for alveoli, capillary inner diameters of ∼15 μm and minimal wall thicknesses of ∼6 μm in plastic prints and 50-100 μm, ∼10 μm and 4 μm for GM10 bioink respectively. We hence could experimentally realize scaffolds that match in scale to human^[74]^ and mouse^[73]^ alveoli. By adjusting design parameters, templates can be personalized, adapted to specific murine or human lung tissue features or adjusted to mimic pathological tissue to for instances investigate fibrotic complications^[75]^. In the future, the direct interface between alveoli walls and capillaries way serve as an organotypic 3D *in vitro* environment to explore direct interactions between alveoli specific pneumocytes and capillary specific endothelial cells.

## 3. Conclusion and Outlook

The high complexity across all scales renders it difficult to fully recapitulate *in vivo* organ and tissue structure in all details encountered *in vitro*. Classic organ-on-a-chip approaches focus on cell biological aspects such as the influence of varying levels of oxygen concentration, corresponding to different zones throughout the human airways^[76],[77]^.

More organotypic microfabrication can help to advance our understanding of 3D cellular dynamics and microtissue physiology in their native context. Our bio-hybrid engineering approach combines microsystems engineering of hydrogel-based 3D geometries and design synthesis. TPS allowed fabrication of both, hydrogel scaffolds with low μm channel diameters, as well as plastic adapters to seamlessly interface standard microfluidic equipment. Using a dip in TPS printing approach enables proteinaceous microscaffolds with arbitrary heights^[37]^ and high precision, due to reduced spherical aberrations during the printing process^[37],[44]^. Other high-resolution imaging techniques such as high resolution serial sectioning electron microscopy^[78]^ or X-ray tomographic phase-contrast imaging for non-destructive 4D imaging of perfused soft microtissues constructs^[7],[79]^ can give detailed insights into the dynamics of 3D printed micro-vasculature models rendering our ultracompact design beneficial.

Further refining cell biological and biophysical properties^[37]^ of the ultracompact ECM hydrogel perfusion chip will enable systematic studies to elucidate the role of context specific tissue microstructures, e.g. in tumor heterogeneity or developmental programs. Tailored photoresin formulations may realize select biodegradability or provide local stimuli for cell specific cell attachment or differentiation under for instances curvotactic influence^[80]–[82]^. Integrating pressure sensors and actuators to the microfluidic circuit can further improve control over flow dynamics and shear stress to optimally recapitulate *in vivo* conditions^[83]^.

The algorithmic design principles used here are well suited to realize biomimetic microtissue models allowing for selective topography variations and artificial randomness. Increasingly detailed 3D imaging data may be incorporated into the algorithmic design process and profoundly improve our capacity to 3D print organotypic tissue. In the future, algorithmically derived print templates will hence not only include geospatial data, but also local print parameters such as laser intensity distributions, to realize e.g. biophysical (stiffness) or biochemical (composition) gradients within complex designs.

The design freedom inherent to additive manufacturing offers other tantalizing avenues for the design and system architectures presented here. These may include algorithmic design synthesis of efficient bioreactors or multimaterial nozzles for bioprinting approaches^[84]^.

## 4. Material and Methods

Unless stated otherwise, all chemicals and cell culture reagents were purchased from Sigma-Aldrich (St. Louis, MO, USA).

### High-Resolution 3D Printing

All adapters and GM10 scaffolds were designed with SolidWorks 2020 (Dassault Systèmes). Designs were exported as STL files and converted to print job instructions using Describe followed by selective exposure performed with the Nanoscribe GT Photonics Professional operating in dip-in mode with an erbium-doped femtosecond laser source and a center wavelength of 780 nm. Power amounted to 150 mW using a 25× (NA 0.8) objective. Adapters were printed using IP-S resin (all Nanoscribe GmbH) while cell scaffolds were printed using GM10 resin. Scaffolds which were not connected to TPS printed adapters were printed within 35 mm glass bottom petri dishes (MatTek) as previously described^[37]^.

### GM10-based resin

Gelatin methacryloyl was synthesized and characterized as previously discussed using gelatin type B (Limed, bovine bone, 232 g Bloom, viscosity: 4.5 mPa s, Gelita, Germany), yielding GM10 (degree of methacrylation:1.07 mmol g^−1^)^[63]^. GM10 stock solution was prepared by dissolving GM10 in PBS. Lithium phenyl-2,4,6-trimethylbenzoylphosphinate (LAP, >95% purity) stock solution was prepared by dissolving LAP in ddH_2_O with a stock concentration of 340 mM. Rose bengal (>95% purity) was dissolved in PBS buffer to yield a stock concentration of 85 mM. The rose bengal stock solution was centrifuged for 5 min at 13 × 103 rpm to sediment undissolved impurities. GM10 stock solution was mixed with LAP and RB stocks resulting in final concentrations of 25 wt% GM10, 68 mM LAP and 0.5 mM RB as previously described^[37]^.

### Microfluidics

Fully assembled adapters containing precision printed 3D protein scaffolds were connected to pressure-based flow control (FlowEZ, 345 mbar, Fluigent) by MicroTight adapters to interface 1/32inch tubing (IDEX Health & Science LLC) with 360 μm OD HPLC capillary tubing (PolyMicro).

### Cell handling

Human mesenchymal stem cells (hMSCs) were cultured in DMEM (BS.FG 0445, Bio&SELL GmbH) supplemented with 1% penicillin-streptomycin (BS.A 2213, Bio&SELL GmbH), 10% fetal calf serum (F7524-500mL, Sigma-Aldrich Chemie) and 1% GlutaMAX (35 050 038, Life Technologies GmbH) under humidified conditions at 37 °C and 5% CO_2_. Cell culture media was exchanged every 2-3 days and cells were split at 80–90% confluency using 0.5% trypsin-EDTA solution (BS.L 2163, Bio&SELL GmbH), centrifuged at 500 rpm for 5 min at room temperature and finally seeded in a ratio of 1:6 in T175 cell culture flasks (83.3912.002, Sartstedt AG und Co.).

### Stereomicroscopic imaging

Adapter assembly and stereomicroscopic images were conducted with a Wild M450 Epimicroscope (Leica Microsystems GmbH).

### Epifluorescence microscopy

Live cell fluorescence imaging of hMSCs was conducted on a AxioObserver.Z1 (Zeiss) using ZEN blue software (Zeiss, 2009) with 10x (NA: 0.3) and 32x (NA: 0.45) objectives.

### Scanning electron microscopy

was used to image 3D prints fabricated in IP-S resin prints. After sputtering approximately 1 nm platinum coated using a Cressington 208HR prints were imaged on a LYRA3 (Tescan Orsay Holding) at 4.0 kV in high vacuum.

### Two photon microscopy

A homebuilt two-photon excited fluorescence microscope was used for diffraction-limited 3D imaging of printed scaffolds under live cell conditions. Compared to linear fluorescence microscopy methods, multiphoton microscopy can achieve higher-resolution and deeper penetration depths while maintaining low photo toxicity. A Nikon Eclipse Ti2 body was modified with custom 3D printed parts to adopt it for a broad variety of multiphoton and nonlinear microscopy techniques. A 100 mW, 95 fs laser pulsed at 80 MHz (FemtoFiber Dichro Design, TOPTICA Photonics AG) with wavelengths of *λ*_1_=1034 nm and *λ*_2_=780 nm was used as excitation source. The two beams were combined by a dichroic mirror (F38-825, AHF Analysentechnik) before they were coupled into a resonant-galvo scanner system (Multiphoton Essential Kit, Thorlabs). A 20x, 0.95 NA water immersion objective lens (CFI Apo MRD77200, Nikon) was used for laser scanning. Fluorescence light from the sample is collected by the same objective lens in epi direction, before it is separated from the illumination beam path by a dichroic mirror (FF825-sDi01, Laser2000) and quantified in a non-descanned configuration with two InGaAsP photomultiplier tubes (Multiphoton Essential Kit, Thorlabs). A dichroic mirror (F76-735, AHF Analysentechnik) in the detection path separates the fluorescence signal in blue (F76-594, AHF Analysentechnik) and red (F32-600A, AHF Analysetechnik) spectral emission ranges. Two multiphoton filters (F39-745, AHF-Analysentechnik) block out scattered excitation light for low background signal and noise.

Image datasets were recorded individually for each excitation wavelength. The sample stage installed in the microscope was synchronized with the image acquisition software by a self-written program. For each z-position, a cumulated image was calculated based on 100 single images. Cumulative images were recorded at subsequent axial positions, leading to an optically sectioned 3D image after image reconstruction. Image analysis was performed using ImageJ Fiji ^[85]^ and self-written Matlab scripts.

To visualize accessible microchannels inside hydrogel 0.5 mg/ml FITC-CM-Dextran (Fluoresceinisothiocyanat-Carboxymethyl–Dextran, average molecular weight 150 kDa, Sigma-Aldrich) in PBS buffer. All PBS inside the petri dish containing 3D printed proteinaceous scaffolds were then replaced with 3 ml of stain solution and imaged after at least 10 min.

### Algorithmic design

Algorithmically derived vascular alveoli print templates were designed in Hyperganic 2.0 software (Hyperganic Group GmbH). Obtained designs are available in STL format through University Stuttgart DaRUS data repository https://doi.org/10.18419/darus-2612.

### Statistical analysis

Proteinaceous channels were printed onto custom adapters and perfused using PBS (n = 3) or cell suspension (n = 2). Scaffolds to determine channel resolution were printed using TPS (n = 3) and imaged using two-photon microscopy. Computational alveoli templates were printed using acrylate-based IP-S resin (n = 3) or proteinaceous GM10-based resin (n = 5).

## Acknowledgements

Funding from the Bavarian State Ministry of Science and the Arts through the “Bavarian Research Institute for Digital Transformation (bidt)” fellowship to AE, the “BayWISS-Kolleg Ressource Efficiency and Materials” and the Bavarian Research Focus “Herstellung und biophysikalische Charakterisierung von dreidimensionalen Geweben (CANTER), as well as the Ministry of Science, Research and Arts of Baden-Württemberg and the University of Stuttgart within the “Leistungszentrum Mass Personalization” (https://www.masspersonalization.de/)and 3R-Netzwerk-BW (https://www.the3rs.uni-tuebingen.de/) is gratefully acknowledged.

We thank Alexander Southan, Jana Grübel and Anastasia Tsianaka (IGVP, University of Stuttgart) for providing the GM10 used in this study, Conny Hasselberg-Christoph (CANTER, Munich University of Applied Sciences) for comprehensive cell culture support, Petra Schwille and Frank Siedler (MPI of Biochemistry) for generous microfabrication lab access and support, Hyperganic Group GmbH including Lin Kayser and Markus Finke for providing early access to Hyperganic software, and Oliver Hayden (Biomedical Electronics, Technical University Munich).

## Author contributions

AE and MH designed research with contributions from HCS and SS. AE performed chip design, microfabrication and characterization. JL and AE implemented parametric design synthesis of vascularized alveoli. TK and TH performed two-photon imaging and data processing. CE performed SEM imaging. MH, HCS and SS supervised research, interpreted data and acquired funding. AE and MH wrote the manuscript with contributions from JL, TK and revisions from HCS and SS. All authors approved the final manuscript.

