## supplement for "Engineering Principles and Algorithmic Design Synthesis for Ultracompact Bio-Hybrid Perfusion Chip"

### This PDF file includes:

Figure S1
Captions for Movies S1 to S3
Captions for Supplemental Design Files

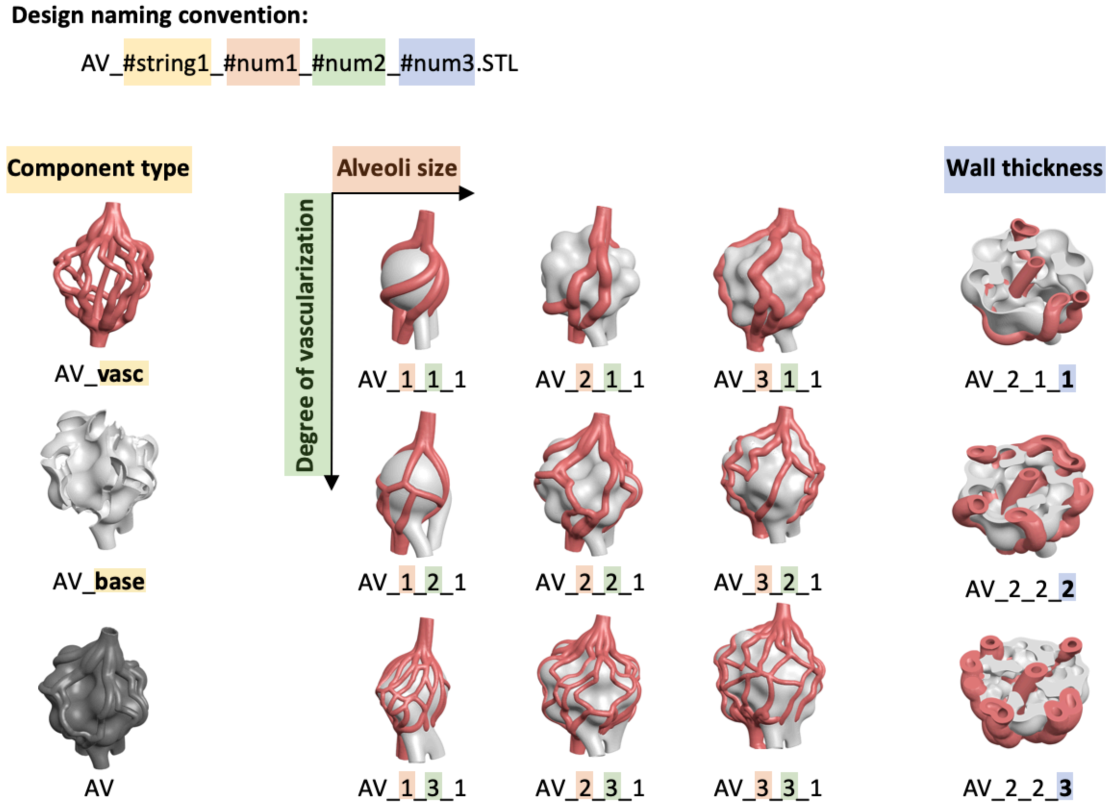


*Figure S1:* Vascularized alveoli designs generated from parametric design synthesis to realize context specific 3D design cues. Implemented variations include component type (#string1), alveoli size (#num1), degree of vascularization (#num2), and wall thickness (#num3) are available from University Stuttgart DaRUS data repository: https://doi.org/10.18419/darus-2612 .

### Movie Captions:

*Movie S1*: Movie of 80 µm diameter GM10 channel perfused with PBS via IP-S printed contact chip (Figure 3).
*Movie S2*: A 3D view and a sliced computer representation of a algorithmically designed alveoli (Figure 7).
*Movie S3*: Animated two-photon microscopy z-stack of an algorithmically designed alveoli printed in GM10 – rose bengal resin (Figure 7E, scale bar 100 µm).

### Designs files:

*Design S1:* Contact Chip Template used for IP-S printing (Figure 1).
*Design S2:* Channel Template used to print from hydrogel GM10 resin (Figure 1).
*Algorithmic Design files:* AV_#string1_#num1_#num2_#num3.STL as annotated in Figure S1 available from https://doi.org/10.18419/darus-2612 .
